# Defining a canonical ligand-binding pocket in the orphan nuclear receptor Nurr1

**DOI:** 10.1101/278440

**Authors:** Ian Mitchelle S. de Vera, Paola Munoz-Tello, Venkatasubramanian Dharmarajan, David P. Marciano, Edna Matta-Camacho, Pankaj Kumar Giri, Jinsai Shang, Travis S. Hughes, Mark Rance, Patrick R. Griffin, Douglas J. Kojetin

**Affiliations:** Department of Integrative Structural and Computational Biology, The Scripps Research Institute, Jupiter, Florida 33458, USA; Department of Molecular Medicine, The Scripps Research Institute, Jupiter, Florida 33458, USA; Skaggs Graduate School of Chemical and Biological Sciences, The Scripps Research Institute, Jupiter, Florida 33458, USA; Department of Molecular Genetics, Biochemistry and Microbiology, University of Cincinnati, Cincinnati, OH 45267, USA; Present address: Department of Pharmacology & Physiology, Saint Louis University, St. Louis, Missouri, 63104, USA; Present address: Department of Biomedical & Pharmaceutical Sciences, The University of Montana, Missoula, Montana 59812, USA

## Abstract

**Nuclear receptor related 1 protein (Nurr1/NR4A2) is an orphan nuclear receptor that is considered to function without a canonical ligand-binding pocket. A crystal structure of the Nurr1 ligand-binding domain (LBD) revealed no physical space in the conserved region where other nuclear receptors with solvent accessible apo-protein ligand-binding pockets bind synthetic and natural ligands. Using solution NMR spectroscopy, hydrogen/deuterium exchange mass spectrometry, and molecular dynamics simulations, we show here that the putative canonical ligand-binding pocket in the Nurr1 LBD is dynamic with high solvent accessibility, exchanges between two or more conformations on the microsecond-to-millisecond timescale, and can expand from the collapsed crystalized conformation to allow binding of unsaturated fatty acids. These findings should stimulate future studies to probe the ligandability and druggability of Nurr1 for both endogenous and synthetic ligands, which could lead to new therapeutics for Nurr1-related diseases, including Parkinson’s disease and schizophrenia.**

## INTRODUCTION

The orphan nuclear receptor transcription factor Nurr1 (NR4A2) is expressed in the embryonic ventral midbrain and is essential for the development and maintenance of dopaminergic neurons (Kadkhodaei et al., 2009; (Zetterstrom et al., 1997). Nurr1 mutations and polymorphism have been found in patients with Parkinson’s disease (Grimes et al., 2006; (Le et al., 2003; (Xu et al., 2002). Post-mortem analysis of human brain tissue from neuropathologically verified cases of Parkinson’s disease revealed downregulation of *NURR1* expression, indicating a role for Nurr1 in the decreased production of dopamine and degeneration of dopaminergic neurons (Decressac et al., 2013). These observations implicate Nurr1 as a therapeutic target for treating Parkinson’s disease, which is supported by studies showing that synthetic Nurr1 agonists show neuroprotective effects in animal models of Parkinson’s disease (Kim et al., 2015; (Kim et al., 2016; (Zhang et al., 2012).

Crystal structures have revealed that endogenous/ natural and synthetic nuclear receptor ligands bind to a canonical ligand-binding pocket within the core of the ligand-binding domain (LBD). However, crystal structures of the LBD of Nurr1 and two related NR4A subclass orphan nuclear receptors, Nur77 (NR4A1) and the *Drosophila* ortholog DHR38 (NR4A4), show no apparent ligand-binding cavity in this same physical space (Baker et al., 2003; (Flaig et al., 2005; (Wang et al., 2003). Instead, their putative canonical ligand-binding pockets are filled with bulky hydrophobic residues. These NR4A crystal structures, coupled with the observation that the NR4As display high cellular activation in the absence of added exogenous ligand, have led to their classification as ligand-independent transcription factors—a classification shared with other orphan nuclear receptors with no apparent crystallized canonical ligand-binding pocket (Gallastegui et al., 2015).

Structural studies have shown that other orphan nuclear receptors with previously unidentified endogenous ligands can undergo large conformational changes to bind ligand. A prominent example involves the NR1D subclass known as the REV-ERBs. Although a crystal structure of ligand-free (apo)-REV-ERBβ revealed a collapsed pocket (Woo et al., 2007), a subsequent crystal structure revealed its putative pocket dramatically expands by 600 Å3 to accommodate binding of the porphyrin heme (Gallastegui et al., 2015; (Pardee et al., 2009), which was identified as an endogenous REV-ERB ligand (Raghuram et al., 2007; (Yin et al., 2007). Using NMR spectroscopy we showed that the ligand-free REV-ERBβ ligand-binding pocket is dynamic (Matta-Camacho et al., 2014), confirming the crystallography observations that its apo-pocket has the ability to expand. Furthermore, NMR studies have revealed dynamic apo-pockets for other non-orphan nuclear receptors with large crystallized pockets that bind endogenous ligands, including PPARγ (Hughes et al., 2012; (Johnson et al., 2000), RXRα (Lu et al., 2006), and VDR (Singarapu et al., 2011).

Of the few synthetic Nurr1 agonists reported to activate Nurr1 transcription (Dubois et al., 2006; (Hintermann et al., 2007), one class of compounds sharing a 4-amino-7-chloroquinoline scaffold, which includes amodiaquine and chloroquine, was shown to bind the Nurr1 LBD using NMR chemical shift footprinting to map the ligand binding epitope (Kim et al., 2015). Furthermore, although no endogenous ligands are known to regulate Nurr1 activity *in vivo*, mass spectrometry-based metabolomics *in vitro* identification studies showed that unsaturated fatty acids present in brain tissue bind to the LBDs of Nur77 (Vinayavekhin and Saghatelian, 2011) and Nurr1 (McFedries, 2014). Using NMR chemical shift footprinting, we showed that one of the most enriched brain unsaturated fatty acids in the metabolomics studies, docosahexaenoic acid (DHA), binds to the Nurr1 LBD (de Vera et al., 2016) with a similar NMR-detected binding epitope as the synthetic Nurr1 agonist amodiaquine (Kim et al., 2015).

A preliminary NMR study of the Nurr1 LBD indicated that residues comprising the putative ligand-binding pocket had shorter *T*_2_ relaxation times, suggesting flexibility or dynamics on the μs-ms timescale (Michiels et al., 2010). We were therefore interested in the structural mechanism that could potentially allow Nurr1 to accommodate a bound natural ligand given the apparent collapsed Nurr1 pocket conformation captured by crystallography. Here, using NMR spectroscopy and hydrogen/deuterium exchange coupled to mass spectrometry (HDX-MS), which are sensitive to protein dynamics in solution, we show that the putative Nurr1 ligand-binding pocket is dynamic, solvent accessible, and exchanges between two or more conformations. Using NMR chemical shift footprinting, we show that the binding epitope for several unsaturated fatty acids identified by metabolomics *in vitro* identification studies to bind Nurr1 (McFedries, 2014) colocalizes with the dynamic putative pocket. Finally, using conventional and accelerated molecular dynamics simulations, we show that the collapsed Nurr1 ligand-binding pocket captured by crystallography can expand to a conformation similar to other fatty acid-bound nuclear receptors.

## Results

### The putative ligand-binding pocket exchanges between two or more conformations in solution

Solution NMR peak lineshape analysis reports on dynamic processes including binding events and conformational changes (Akke, 2002; (Kleckner and Foster, 2011). When we compared NMR peak intensities and lineshapes of backbone amides and side-chain methyl groups in the Nurr1 LBD (**Figure 1A–C**), residues within structural regions comprising the putative ligand-binding pocket showed reduced NMR peak intensities and broad lineshapes compared to residues in regions remote from the pocket. These features are indicative of dynamics on the microsecond-to-millisecond (μs–ms) timescale, such as a exchange between two or more structural conformations in solution. To confirm this directly, we performed constant-time Carr-Purcell-Meiboom-Gill relaxation dispersion (CT-CPMG RD) experiments that directly probe chemical exchange (*R*_ex_) processes on the μs-ms timescale. Elevated *R*_ex_ rates were observed for 28 residues in the Nurr1 LBD (**Figure 1D**), which primarily map to the same structural regions that show reduced NMR peak intensities. We also performed ^19^F-NMR by covalently labeling the five cysteine residues present in the Nurr1 LBD with 3-bromo-1,1,1-trifluoroacetone (BTFA), a small molecule compound containing an NMR-observable trifluoromethyl group (–CF_3_) that is highly sensitive to environment (Kitevski-LeBlanc and Prosser, 2012). 19FNMR signals corresponding to BTFA labels attached to C465 and C475, which are located within the putative pocket, show much lower intensity relative to C534 and C566 located outside of the pocket (**Figure 1E, Supplementary Figure S1**). The peak pattern for C465 in particular is complex, with several visible peaks with very broad lineshapes. When plotted onto the crystal structure of the apo-Nurr1 LBD (Wang et al., 2003), the NMR data showing complex and/or broad NMR lineshapes and elevated *R*_ex_ rates all map to the putative pocket (**Figure 1F**). Combined, these data show that the putative apo-Nurr1 ligand-binding pocket exchanges between two or more conformations in solution on the μs-ms timescale.

**Figure 1.**
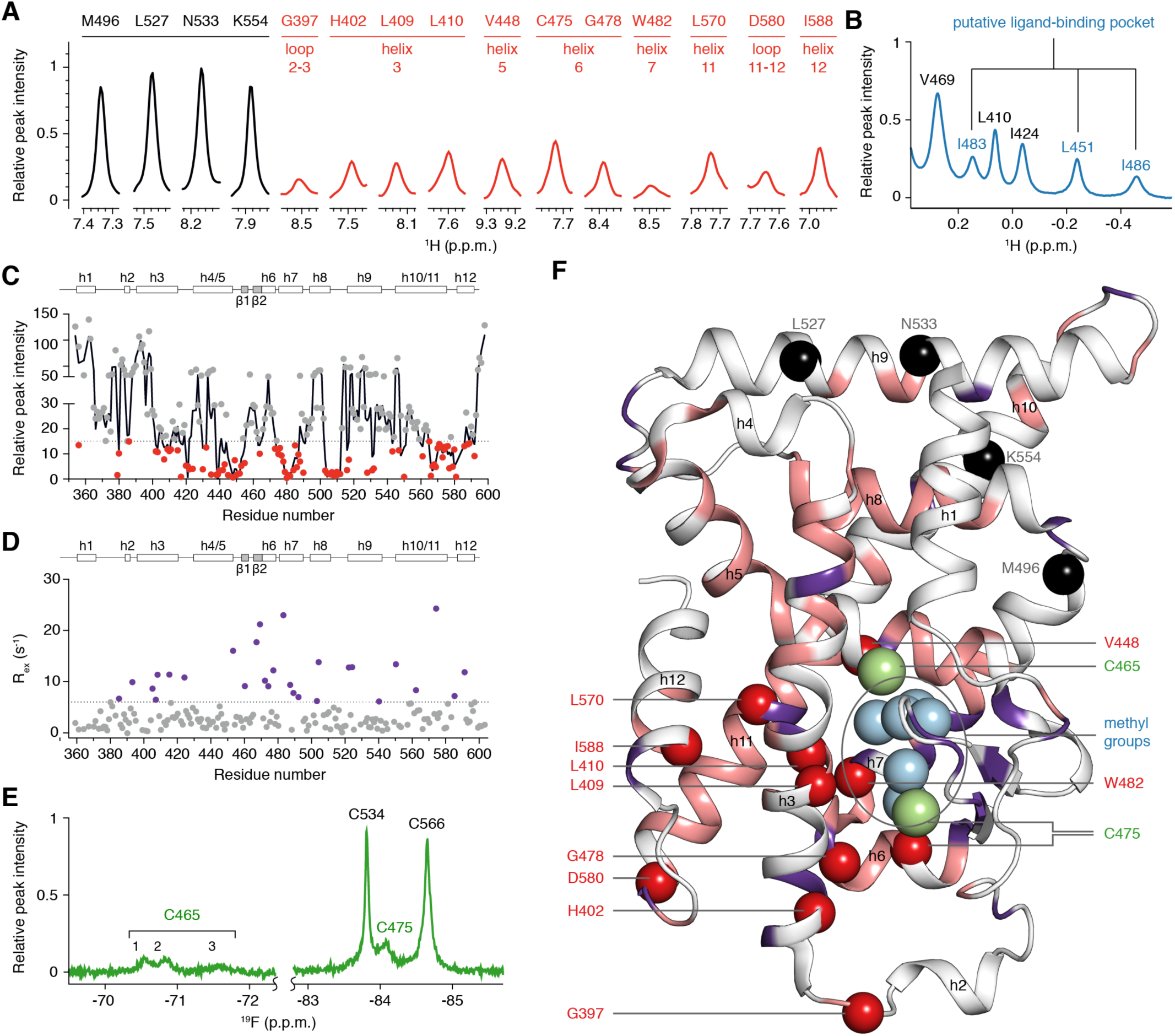
NMR detected μs-ms timescale dynamics within the putative Nurr1 ligand-binding pocket. (**A–C**) NMR lineshape analysis of the apo-Nurr1 LBD in (**A**) extracted 1D slices of 2D [^1^H,^15^N]-TROSY-HSQC NMR data of select backbone amide groups (broad resonances colored and labeled red), (**B**) 1D [^1^H]-NMR data of methyl groups (broad resonances labeled blue), and (**C**) peak intensities of 3D TROSY-HNCO NMR data (residues with peak intensities < 15 colored red; black line represents a two residue running average). (**D**) Chemical exchange (R_ex_) values measured at 18.8 T. Residues with Rex values > 8 s^−1^ are colored purple. (**E**) 1D [^19^F]-NMR data of BTFA-labeled cysteine residues of apo-Nurr1 LBD (broad resonances labeled green). (**F**) NMR data mapped onto the crystal structure of apo-Nurr1 LBD (PDB 1OVL; chain B); backbone amide groups with sharp (black) or broad (red) backbone amide groups in (**A**), broad methyl groups (light blue) in (**B**), HNCO peak intensities < 15 (light pink) in (**C**), R_ex_ values > 8 s^−1^ (purple) in (**D**), and broad BTFA-cysteine resonances (light green) in (**E**).

### The putative pocket has high solvent accessibility

Hydrogen/deuterium exchange analysis coupled to mass spectrometry (HDX-MS) is a solution-based structural method that is sensitive to conformational dynamics, including the detection of protein “breathing motions” and energetically excited conformational states (Konermann et al., 2011). HDX coupled to mass spectrometry (MS) has shown apo-pockets of nuclear receptors have high solvent accessibility (Bruning et al., 2007; (Musille et al., 2012; (Zhang et al., 2010). The crystal structure of the Nurr1 LBD, which shows no apparent pocket volume (Wang et al., 2003), suggests the putative ligand-binding pocket would likely have no apparent solvent accessibility. However, our NMR data above indicates the putative pocket is dynamic and may exchange among other conformations not captured in the crystal structure. We therefore performed HDX-MS to assess the relative solvent exposure of the putative pocket relative to other solvent exposed structural regions (**Figure 2**).

**Figure 2.**
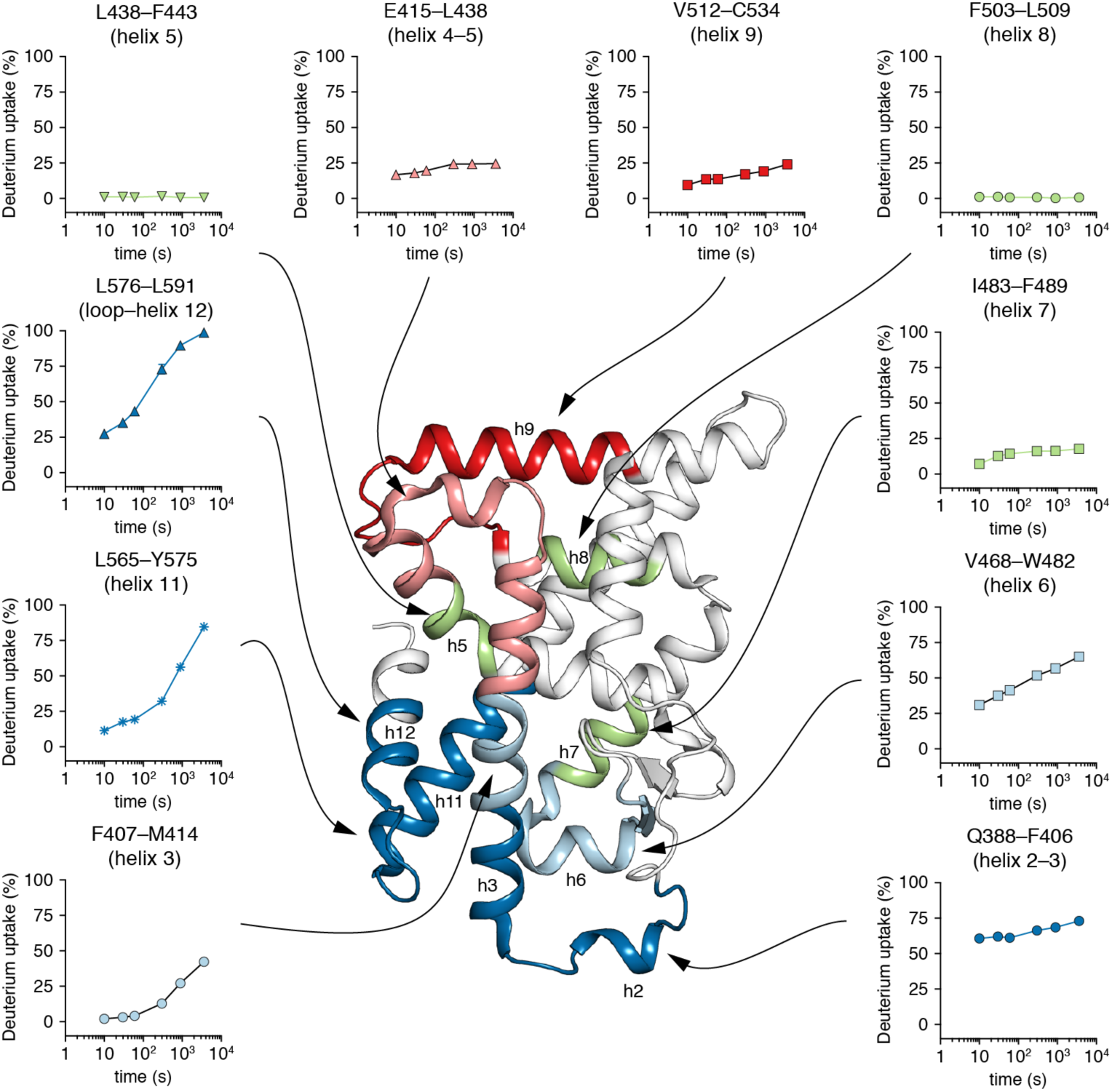
Solvent accessibility of Nurr1 LBD probed by HDX-MS. Deuterium uptake curves show that peptides within the putative ligand-binding pocket have high solvent exchange (blue) relative to the low-to-moderate exchange observed for peptides within the core of the Nurr1 LBD (green) and other solvent exposed peptides on the surface of the Nurr1 LBD (red). Data represent the mean and s.d. of three experimental replicates. Analyzed peptide fragments are colored and displayed on the Nurr1 LBD crystal structure (PDB 1OVL; chain B) using the same colors in the deuterium uptake plots.

Peptides corresponding to residues on the surface of the Nurr1 LBD that would be predicted to have a large exposure to solvent, such as helix 4–5 (E415–L438) and helix 9 (V512–C534), showed slight to moderate exchange with solvent. This indicates these regions are structurally rigid although they could be expected to have high solvent exposure based on their locations and surface accessibility in the crystal structure. Other peptides in structural regions within the core of the LBD similarly show either negligible exchange with solvent, including helix 5 (L438–F443) and helix 8 (F503–L509) or moderate exchange in the case of helix 7 (I483–F489). In contrast, peptides within the putative pocket region show a large degree of exchange with solvent suggesting structural flexibility and high solvent accessibility. This includes peptides within helix 2–3 and the connecting loop (Q388–F406), helix 3 (F407–M414), helix 6 (V468–W482), helix 11 (L565–Y575), and helix 12 and the preceding loop (L576–L591). These highly solvent accessible peptides colocalize to structural regions that our NMR studies showed have μs-ms timescale dynamics.

### Crystallography likely captured a collapsed pocket

Atomic displacement factors, also called *B*-factors or temperature factors, report on thermal motion or vibration and static disorder in structural models derived from x-ray crystallography data. High *B*-factor values are typically observed in the ligand-binding pockets of nuclear receptors known to bind natural or endogenous ligands such as PPARγ (Nagy and Schwabe, 2004), which is supported by dynamic data derived from NMR and HDX-MS studies (Bruning et al., 2007; (Hughes et al., 2012; (Johnson et al., 2000). In the six Nurr1 LBD molecules within the crystallography asymmetric unit (Wang et al., 2003), higher than average *B*-factor values (**Figure 3A**) are observed for residues comprising the putative ligand entry-exit surface formed at the base of LBD where helix 3 and 11 meet with the helix 6–7 loop, as well as the helix 2–3 loop (Edman et al., 2015; (Grebner et al., 2017). Unfortunately, structure factors were not deposited for the Nurr1 LBD crystal structure, which limits further interpretation of the electron density of this structure. However, electron density for the helix 2–3 loop within the region of the putative pocket was apparently poor or absent because this loop was not modeled in four of six molecules in the asymmetric unit, which could imply flexibility within this region. Consistent with the crystallographic *B*-factor analysis, steady-state [^1^H,^15^N]-heteronuclear nuclear Overhauser effect (hnNOE) NMR analysis of the apo-Nurr1 LBD (**Figure 3B**) revealed large amplitude mobility (hnNOE values < 0.65) in the helix 1–3 loop in addition to other solvent exposed loop regions (**Figure 3C**). The hnNOE NMR experiment reports on ps-ns timescale motions whereby residues with hnNOE values near 1 are highly restricted in motion, and residues with low hnNOE values have increased flexibility (Kleckner and Foster, 2011). These observations, when considered with our NMR and HDXMS data, indicate that the putative pocket captured by crystallography may represent a collapsed conformation, but in solution the pocket is solvent accessible and exchanges between two or more conformations on the μs-ms timescale.

**Figure 3.**
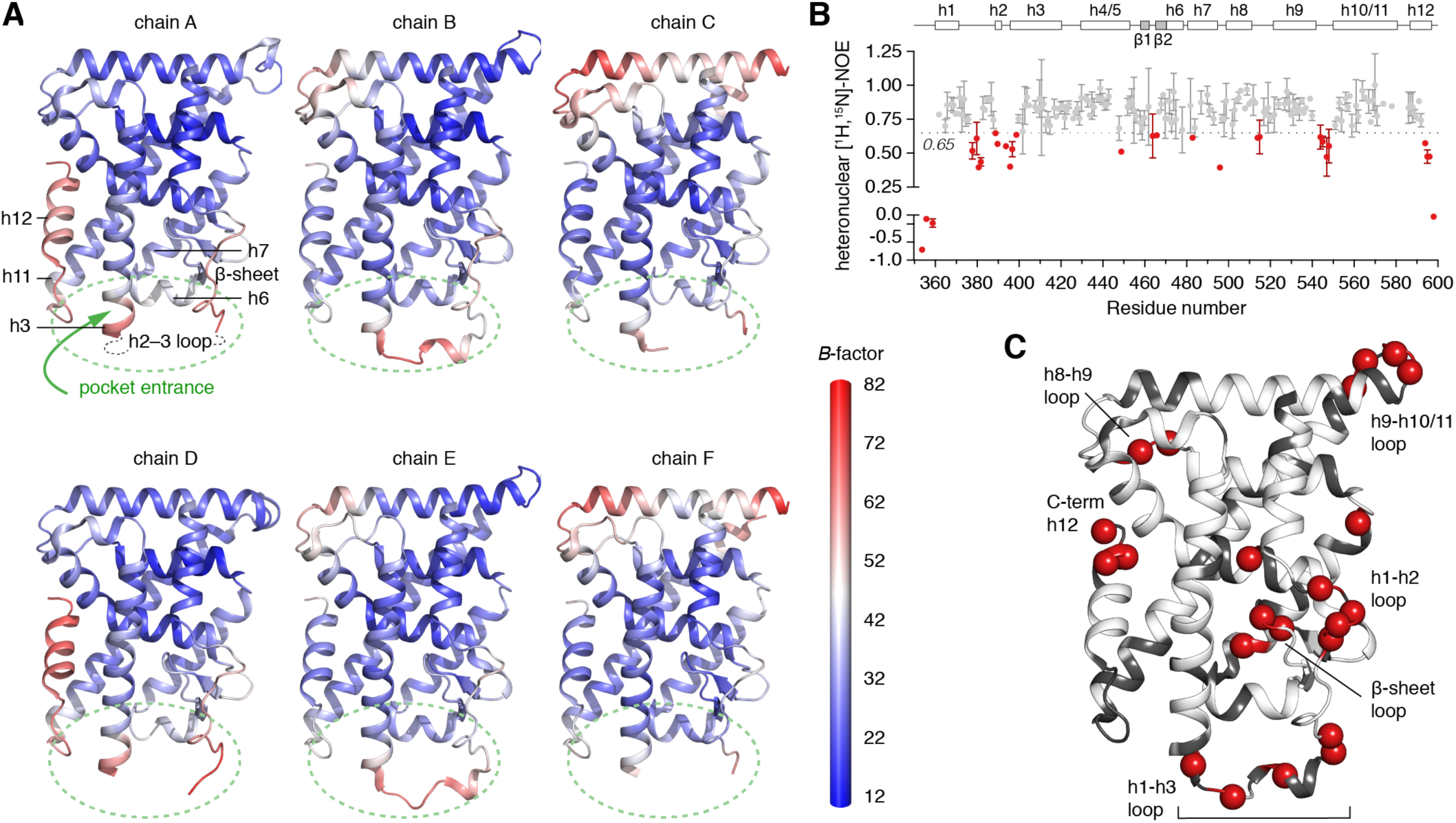
Crystallographic *B*-factors confirm the putative pocket entrance is dynamic. (**A**) *B*-factor values colored from blue (low) to red (high) in chains A–F of the apo-Nurr1 LBD crystal structure (PDB 1OVL). Helical regions are labeled (e.g., helix 12 = h12) and the pocket entrance surface is marked with a green dotted circle. (**B**) Apo-Nurr1 LBD steady-state [^1^H,^15^N]-heteronuclear nuclear Overhauser effect (hnNOE) values measured at 18.8 T. Data represent the mean and s.d. of two individual measurements. (**C**) Residues with hnNOE values < 0.65 in (red spheres), indicative of increased flexibility, mapped onto the crystal structure of apo-Nurr1 LBD crystal structure (PDB 1OVL; chain B).

### Unsaturated fatty acids bind to the dynamic putative pocket

Mass spectrometry-based *in vitro* metabolomics studies identified unsaturated fatty acids present in metabolic extracts from mouse brain that bind to the LBDs of Nurr1 (McFedries, 2014) and Nur77 (Vinayavekhin and Saghatelian, 2011), an orphan nuclear receptor evolutionarily related to Nurr1. Using NMR structural footprinting analysis, we demonstrated that the binding epitope of one of the most enriched ligands identified in the metabolomics studies, docosahexaenoic acid (DHA; C22:6), maps to the putative Nurr1 ligand-binding pocket (de Vera et al., 2016). Several other Nurr1-binding unsaturated fatty acids were identified in the *in vitro* brain metabolite identification studies, including arachidonic acid (AA: C20:4), linoleic acid (LA; C18:2), and oleic acid (OA; C18:1). We used tryptophan fluorescence spectroscopy to validate binding of these unsaturated fatty acids (**Figure 4A**), which provided K_d_ values of 50 ± 5 μM (AA), 101 ± 13 μM (LA), and 59 ± 15 μM (OA). We performed NMR structural footprinting to determine if the binding epitopes of these ligands are similar to DHA (**Figure 4B**). NMR peaks with the largest changes in chemical shift and decrease in peak intensities map to the putative pocket (**Figure 4C**). This binding epitope is similar to NMR detected ligand binding epitopes of Nurr1 binding to DHA (de Vera et al., 2016) and the antimalarial drug amodiaquine (Kim et al., 2015) within the putative pocket where the N-terminus of helix 3 meets the helix 6–7 loop and N-terminus of helix 11, as well as where helix 3 and helix 11 meet helix 5 and helix 12. Furthermore, the Nurr1 ligand binding epitopes are similar to the NMR detected epitopes of ligand binding to PPARγ (Hughes et al., 2012; (Hughes et al., 2014; (Hughes et al., 2016; (Johnson et al., 2000; (Marciano et al., 2015), RXRα (Kojetin et al., 2015; (Lu et al., 2006; (Lu et al., 2009), REV-ERBβ (Matta-Camacho et al., 2014), and VDR (Singarapu et al., 2011). Taken together with our dynamic analyses above, these data indicate the putative pocket can expand from the collapsed crystallized conformation to bind ligand.

**Figure 4.**
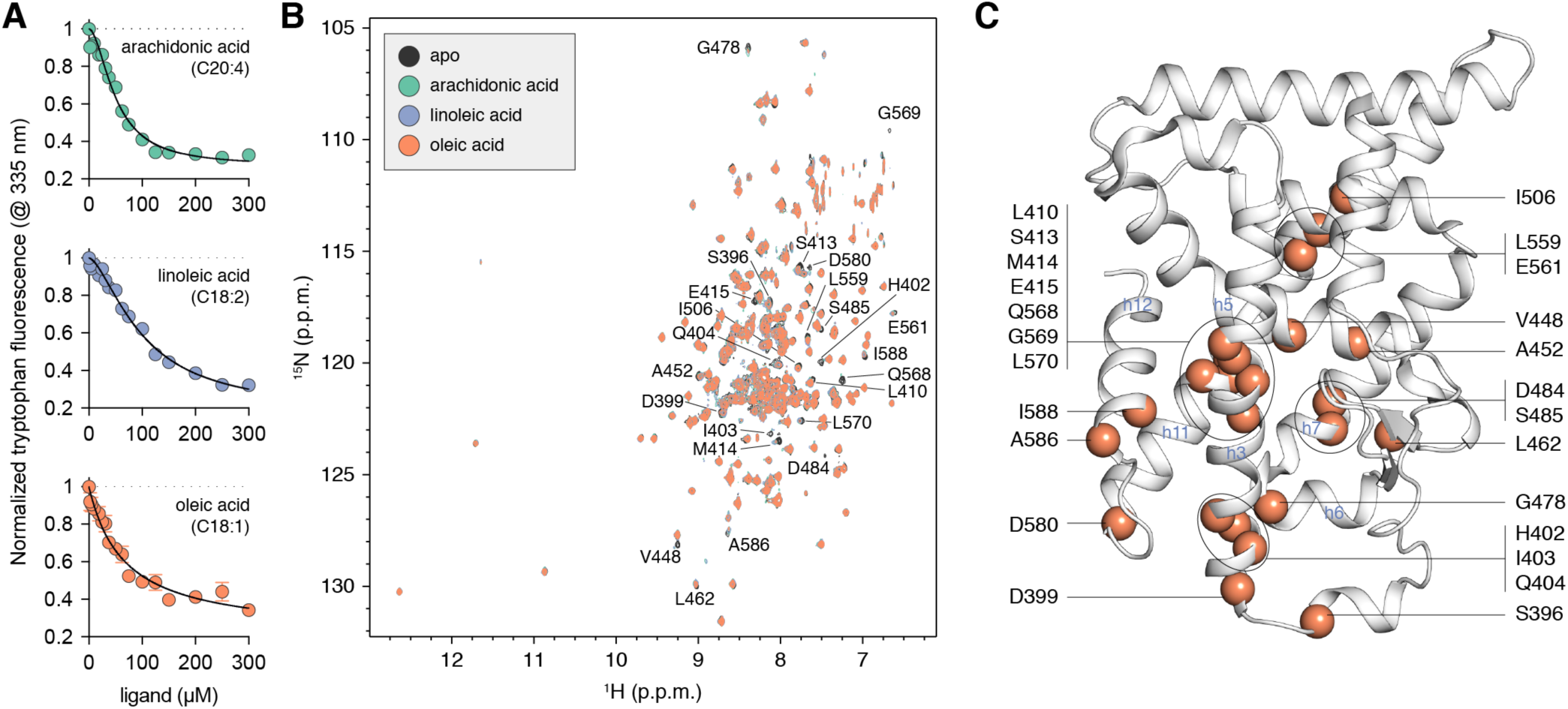
NMR footprinting maps the binding epitope of unsaturated fatty acids to the putative pocket. (**A**) Nurr1 tryptophan fluorescence binding assay for unsaturated fatty acids. (**B**) [^1^H,^15^N]-TROSY-HSQC NMR spectra of Nurr1 LBD in the absence or presence of 1.5 molar equivalent of the indicated unsaturated fatty acids. Residues showing the most significant perturbations are labeled and displayed in (**C**) on the apo-Nurr1 LBD crystal structure (PDB 1OVL; chain B) as orange spheres.

### The putative pocket can expand to a conformation similar to other fatty acid-bound nuclear receptors

We superimposed the apo-Nurr1 LBD crystal structure (Wang et al., 2003) with several crystal structures of nuclear receptor LBDs bound to fatty acids, including PPARγ bound to DHA (Itoh et al., 2008), RXRα bound to DHA or pentadecanoic acid (Egea et al., 2002; (Xu et al., 2004), and HNF4α bound to myristic or palmitic acid (Duda et al., 2004; (Rha et al., 2009; (Wisely et al., 2002), to determine the structural rearrangements that would need to occur in order for the collapsed crystallized apo-Nurr1 ligand-binding pocket conformation to expand (**Figure 5A, Supplementary Figure S2**). The conformation of the loop connecting helix 6 and 7 in the apo-Nurr1 LBD, which is tucked into the putative pocket in close proximity to helix 3, stands out among the other fatty acid-bound nuclear receptor LBD structures, which show an open conformation where the helix 6–7 loop is further away from helix 3. Notably, the helix 6–7 loop region is a critical structural mediator of the ligand entry-exit site within the nuclear receptor LBD (Edman et al., 2015; (Grebner et al., 2017). In our Nurr1 NMR studies, residues in and near the helix 6–7 loop show very broad NMR linewidths and elevated Rex values, indicating exchange between two or more conformations on the μs-ms timescale. Furthermore, residues within this region show NMR chemical shift perturbations upon binding unsaturated fatty acids, revealing this dynamic region is also structurally affected by ligand binding.

**Figure 5.**
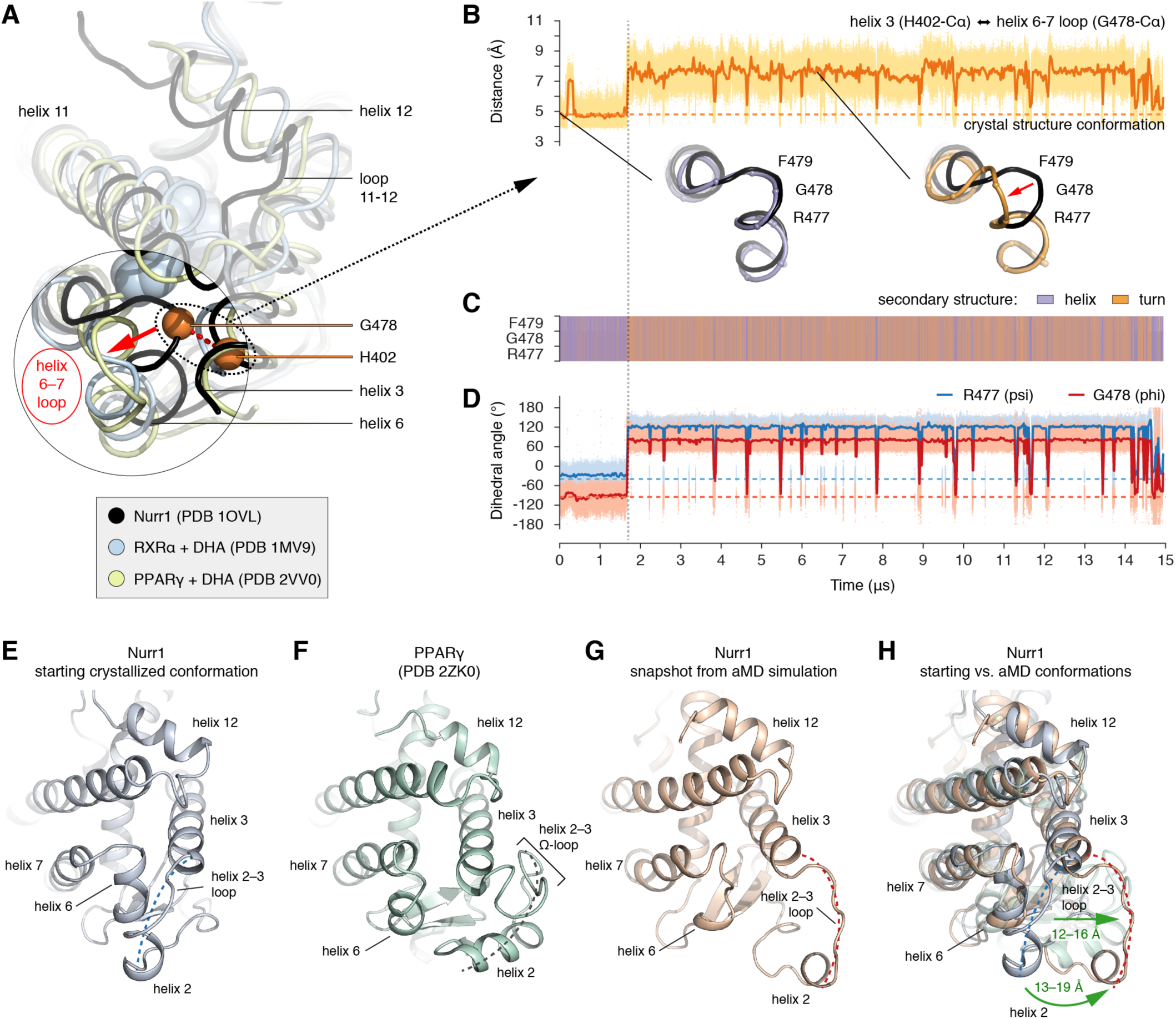
The putative Nurr1 ligand-binding pocket expands in molecular dynamics simulations to conformations similar to other ligand-binding nuclear receptors. (**A**) Structural superposition of the LBDs of apo-Nurr1 (PDB 1OVL) with DHA-bound RXRα (PDB 1MV9) and DHA-bound PPARγ (PDB 2VV0). C atoms of H402 (helix 3) and G478 (loop 6-7) are shown as orange spheres; the distance between this atom pair was monitored in the simulations. (**B-D**) Conventional MD simulations of apo-Nurr1 reveals that (**B**) loop 6-7 moves away from helix 3 (red arrow) during the simulation resulting in an open pocket conformation (purple vs. orange; vs. PDB 1OVL in black), as well as a change in (**C**) secondary structure from helix to turn and (**D**) backbone dihedral angles. Starting conformations are noted by dotted lines, and running averages of the MD data are shown as solid colored lines. (**E–H**) Accelerated MD (aMD) simulations show that the Nurr1 pocket entry-exit site, which (**E**) crystallized in a closed conformation compared to (**F**) the open accessible pocket conformation in PPARγ, (**G,H**) can undergo a large shift in the location of the helix 2/helix 2–3 loop to an open conformation that is similar to PPARγ. Dotted lines note the locations of the helix 2/helix 2–3 loop regions.

The superposition and NMR analyses indicate the dynamic helix 6–7 loop in the apo-Nurr1 LBD would need to change from the closed crystallized conformation to an open conformation similar to crystal structures of fatty acid-bound nuclear receptor LBDs to allow the pocket to expand and bind ligand. To explore this possibility, we performed conventional molecular dynamics (MD) simulations, including 9 independent simulation runs ranging independently from 4.5–15 μs in length. In 8 of the 9 simulations, the helix 6–7 loop moved and extended away from the crystallized starting conformation towards the open conformation observed in the fatty acid-bound nuclear LBD crystal structures (**Figure 5B, Supplementary Figure S3**). This conformational change is associated with a change in secondary structure from a helix in the starting crystallized conformation to a turn for residues within the helix 6–7 loop region (**Figure 5C,D**). This is consistent with the other fatty acid-bound nuclear receptor LBD crystal structures where the helix 6–7 loop adopts a turn type of secondary structure, not a helix as in the Nurr1 crystal structure.

The observation that the helix 6–7 loop changes to and remains in an open conformation in 8 out of 9 simulations indicates that the crystallization conditions may have selected for a collapsed or closed pocket. Furthermore, the crystallized conformation of the apo-Nurr1 helix 2/helix 2–3 loop surface is unique compared to most nuclear receptor LBD crystal structures, as it tucks against helix 6 forming a “clamp” over the pocket entry-exit surface and would prevent ligand access to the pocket (**Figure 5E**). In contrast, for example, the PPARγ helix 2/helix 2–3 loop surface, which is also called the Ω-loop, adopts an open conformation allowing ligands to access the pocket (**Figure 5F**). As mentioned previously, structure factors were not deposited for the apo-Nurr1 crystal structure (Wang et al., 2003). Although the helix 2/helix 2–3 loop surface was modeled into two of the six chains in the apo-Nurr1 LBD crystal structure, the *B*-factor values for this region, which are the only assessment to the structural quality of the loop conformation, are relatively high. Notably, our NMR data show that the pocket and the entry-exit surface are dynamic and exchange between two or more conformations on the μs-ms timescale.

Because conformational changes on the μs-ms timescale are not realistically accessible by conventional MD, we turned to accelerated molecular dynamics (aMD), which allow sampling of conformational changes that occur on the ms timescale and exploration of conformations beyond the energy basin localized around crystallized conformations, which can remain trapped in conventional MD (Pierce et al., 2012). Indeed, aMD simulations revealed an open helix 2/helix 2–3 loop surface conformation (**Figure 5G**), representing a large conformational change compared to the crystallized closed conformation (**Figure 5H**) that is similar in location to the PPARγ Ω-loop (**Figure 5F**). This conformational change increases the solvent accessibility of the surface where helix 3 meets the helix 6–7 loop, and is consistent with our NMR footpriting data showing the binding epitope of unsaturated fatty acids for the same region. These data, which complement the solution NMR and HDX-MS dynamical studies, further indicate that the crystallized Nurr1 pocket conformation is collapsed, but can change to an open and solvent-accessible conformation similar to the LBDs of other ligand-binding nuclear receptors.

## Discussion

Crystallography has revealed that nuclear receptor ligand-binding pockets can vary in size from essentially no pocket volume (i.e., the NR4As) up to 1,600 Å3 in volume (Gallastegui et al., 2015). Although the ligand-binding pockets of nuclear receptors are understood to be conformationally flexible (Nagy and Schwabe, 2004) and adaptable to expand beyond previously crystallized ligand-bound conformations (Molnar et al., 2006; (Suino-Powell et al., 2008; (Togashi et al., 2005), the absence of pocket volumes in crystal structures of Nurr1 and a small cohort of other orphan nuclear receptors including other NR4As, COUP-TFII, DAX, PNR, SHP, and TR4 has led to the conclusion that these receptors have noncanonical, ligand-independent regulatory mechanisms (Gallastegui et al., 2015).

In contrast to these structural observations from static crystal structures, here we employed structural methods capable of detecting protein motions, or dynamics, in solution. Our NMR data show that putative Nurr1 ligand-binding pocket dynamically exchanges between two or more conformations on the μs-ms timescale. This is apparent from the broad lineshapes in 1D and 2D NMR data, the elevated *R*_ex_ values in the CT-CPMG RD experiments, and the appearance of multiple and/or broad ^19^F NMR peaks for cysteine residues within the pocket covalently tagged with BTFA. Using HDX-MS, we found that the putative pocket has high solvent accessibility, greater than other solvent exposed regions of the Nurr1 LBD. Our NMR footprinting data also identified the putative Nurr1 pocket as the binding epitope for unsaturated fatty acids present in metabolite extracts from mouse brain that were pulled down in mass spectrometry metabolomics *in vitro* identification studies. This NMR-detected fatty acid pocket binding epitope is similar to the NMR-detected binding epitope for the synthetic ligand amodiaquine (Kim et al., 2015), indicating that natural and synthetic ligands likely bind to the same pocket in Nurr1. Finally, using molecular dynamics simulations we found that regions of the Nurr1 LBD that in the crystallized conformation would prevent ligand access to the pocket can physically change to a conformation similar to other ligand-bound nuclear receptors, which could allow a ligand to access the Nurr1 pocket. It was previously posited that a ligand binding event could potentially induce a conformational change that would open up a pocket for orphan nuclear receptors lacking a canonical crystallized pocket, such as Nurr1 and Nur77 (Xu and Li, 2008). Thus, it is possible that unsaturated fatty acids either bind to an open pocket conformation in solution, which could be one of the two or more conformations detected by NMR in solution; or induce a conformational change that expands the pocket after an initial encounter complex binding event.

In our NMR dynamical analyses, which overall revealed the putative pocket to be dynamic on the μs-ms timescale, NMR peaks corresponding to residues in the helix 6–7 loop loop were particularly broad, indicating significant μs-ms motion. This region has been implicated as a critical regulatory element in the ligand binding pathway of nuclear receptors (Edman et al., 2015; (Grebner et al., 2017). Furthermore, the helix 2/helix 2–3 loop surface, which is also implicated in the ligand binding pathway of nuclear receptors (Genest et al., 2008; (Martinez et al., 2005), showed lower than average hnNOE values, indicating significant large amplitude motion on the ps-ns timescale. This region in some chains of the crystal structure was modeled as a helix that forms a lid or cap over the pocket entry site. However, analysis of the NMR chemical shift values for Nurr1 residues within this region suggests a lack of secondary structure, unlike the helix modeled into the crystal structure, in addition to the lower than average hnNOE values. Our molecular dynamics simulations showed that both of these regions could change conformation, which could allow ligand access to the Nurr1 pocket. Furthermore, decreased hnNOEs were also observed for the helix 2/helix 2–3 loop the RXRα LBD, and in our past NMR studies on PPARγ we did not observe NMR peaks for most residues in the helix 2–3 loop (also called the Ω-loop) (Hughes et al., 2012), suggesting that this region may generally be conformationally flexible in nuclear receptors. Moreover, the helix 2/helix 2–3 loop region preceding helix 3 is the most variable portion among the 48 human nuclear receptors and thus could have different regulatory roles in providing access to the pocket. Furthermore, NMR resonances for residues on helix 12 in the Nurr1 LBD are observed, whereas helix 12 resonances are broad or missing in the apo-forms of PPARγ (Hughes et al., 2012; (Johnson et al., 2000) and RXRα (Lu et al., 2006). Structurally this could be caused by the polyproline hinge preceding helix 12 in Nurr1, which is also present in Nur77. This polyproline region could afford stability of helix 12 compared to the more flexible hinges in PPARγ and RXRα that only contain one proline residue, which may afford the high cellular activation of Nurr1.

Our work here indicates the Nurr1 pocket conformation captured by crystallography represents a low energy collapsed conformation. However, our 2D NMR studies showed that the pocket exchanges between two or more conformations on the μs-ms timescale. Furthermore, our ^19^F NMR studies show that the pocket populates two or three low energy conformations, one of which could correspond to the crystallized conformation. As mentioned above, a similar situation was observed in structural studies of the REV-ERBβ LBD, where NMR revealed the apo-REV-ERBβ pocket is dynamic on the μs-ms timescale (Matta-Camacho et al., 2014), and crystal structures revealed a dramatic expansion of the collapsed REV-ERB ligand-binding pocket to enable binding of heme (Pardee et al., 2009; (Woo et al., 2007). Crystallography studies on estrogen-related receptor (ERR) also suggest considerable pocket adaptability, as the pocket volumes in ERR crystal structures span 40–1020 Å3 (Gallastegui et al., 2015). In the case of ERRα, which showed a 100 Å3 apo-pocket (Kallen et al., 2004), an expansion would be needed in order to accommodate binding of the recently discovered endogenous ERRα ligand, cholesterol (Wei et al., 2016), which has a molecular volume of ~440 Å3. In another dramatic example, crystal structures revealed that an analog of glucocorticoid receptor (GR) ligand dexamethasone, where the conserved 3-ketone moiety was replaced with a phenylpyrazole group, effectively doubled the pocket volume, which expanded from 540 Å3 in the dexamethasone-bound structure to 1,070 Å3 (Suino-Powell et al., 2008). Expansion of ligand-binding pockets has also been observed in a Ras GTPase, where an NMR fragment screen identified hits that bind KRas and crystal structures of several of the hits revealed that larger fragments could expand a surface pocket binding site (Maurer et al., 2012).

*In vitro* mass spectrometry-based metabolomics studies combined with cellular assays have successfully identified endogenous ligands for PPARγ (Dozsa et al., 2014; (Kim et al., 2011; (Xu et al., 1999) and ERRα (Wei et al., 2016) that regulate their cellular activities. Although NMR and/or mass spectrometry-based metabolomics *in vitro* identification studies revealed that unsaturated fatty acids such as DHA bind to LBDs of Nurr1 (de Vera et al., 2016; (McFedries, 2014) and the related NR4A receptor Nur77 (Vinayavekhin and Saghatelian, 2011). DHA did not cause large transcriptional effects in cell-based Nurr1 reporter assay (de Vera et al., 2016). It is possible that DHA and other unsaturated fatty acids could be exchangeable structural ligands as we previously demonstrated for DHA (de Vera et al., 2016) and consequently do not significantly affect Nurr1 activity, or that other putative natural ligands present in cell culture media or produced within the cells masks the activity of exogenously added DHA. However, our work here shows that natural ligands (i.e., unsaturated fatty acids) can indeed bind to Nurr1 via a canonical ligand-binding pocket. This finding should inspire future work to seek out endogenous ligands that could regulate the physiological activity of Nurr1 and stimulate the development synthetic Nurr1 ligands.

## Acknowledgements

This work was supported in part by National Institutes of Health (NIH) grants R01GM114420 (DJK), F32DK108442 (RB), R00DK103116 (TH); American Heart Association (AHA) fellowship award 16-POST27780018 (RB); National Science Foundation (NSF) funding to the Summer Undergraduate Research Fellows (SURF) program at The Scripps Research Institute [Grant 1659594]; and the Academic Year Research Internship for Undergraduates (AYRIU) program at The Scripps Research Institute. A portion of this work was performed at the National High Magnetic Field Laboratory (NHMFL/MagLab), which is supported by National Science Foundation (NSF) Cooperative Agreement No. DMR-1157490 and the State of Florida; we thank Mr. Ashley Blue at the NHMFL for assistance with NMR experiments. NMR data presented herein were collected in part at the City University of New York Advanced Science Research Center (CUNY ASRC) Biomolecular NMR Facility; we thank Dr. James Aramini at the ASRC for assistance with NMR experiments.

## Author contributions

I.M.S.dV., P.M.-T., E.M.-C., P.G., J.S., and D.J.K prepared samples, performed NMR analysis and/or tryptophan fluorescence. V.D. and D.P.M. performed HDX-MS experiments supervised by P.R.G. M.R., T.S.H., and D.J.K. performed the molecular dynamics simulations. D.J.K. conceived the experiments and wrote the manuscript with assistance from I.M.S.dV. and P.M.-T. and input from all authors.

## Methods

### Reagents, protein expression/purification, and ligands

Nurr1 LBD (NR4A2; residues 353–598) protein was expressed in *Escherichia coli* BL21(DE3) cells (Life Technologies) using a pET-46 as a tobacco etch virus (TEV) protease-cleavable N-terminal His-Tag fusion protein and purified as previously described (de Vera et al., 2016) in either autoinduction media or M9 media supplemented with ^15^NH_4_Cl (Cambridge Isotope Labs, Inc.). Protein in wash buffer (50 mM Tris pH 7.4, 500 mM NaCl, 7.5 mM imidazole, 5 mM TCEP) was eluted against a 500 mM imidazole gradient through a Ni-NTA column, subsequently incubated with TEV protease at 4 °C overnight to cleave the hexahistidine tag, and loaded anew onto the Ni-NTA column. The flow through containing purified protein was collected, buffer-exchanged into NMR buffer (20 mM KPO_4_ pH 7.4, 50 mM KCl, 5 mM TCEP and 0.5 mM EDTA), and stored at −80 °C. Reagents were obtained from Fisher Scientific unless otherwise indicated. TCEP was obtained from Gold Biotechnology, Inc. Arachidonic acid, linoleic acid, and oleic acid were obtained from Sigma and dissolved in ethanol or ethanol-d6 for NMR experiments.

### NMR spectroscopy

Protein NMR data were acquired on a Bruker 700 MHz NMR spectrometer equipped with a QCI cryoprobe at 298 K. We previously validated (de Vera et al., 2016) the reported Nurr1 LBD NMR chemical shift assignments (Michiels et al., 2010) using 2D [^1^H,^15^N]-TROSY; 3D TROSY-based HNCO, HNCA, HN(CO)CA, HN(CA)CB, and HN(COCA)CB); and 3D ^15^N-NOESYHSQC. For ligand titrations, 2D [^1^H,^15^N]-TROSY were acquired at 298 K using 100 μM ^15^N-labeled Nurr1 LBD in the absence or presence of 150 μM unsaturated fatty acid in NMR buffer containing 10% D_2_O (Sigma). ^15^NCT-CPMG RD and hnNOE experiments were acquired on a Bruker 800 MHz NMR spectrometer equipped with a TCI cryoprobe at 298 K. ^19^F NMR data were acquired on a Bruker 400 MHz NMR spectrometer equipped with a BBFO probe at 298 K using 100 μM Nurr1 LBD and Cys-to-Ala mutants (C465A, C475A, C505A, C534A, and C566A) incubated with 8 mM of 3-bromo-1,1,1,-trifluoroacetone (BTFA; Sigma) for 12 h at 4 °C. Excess BTFA was removed by buffer exchange before concentrating the BTFA-labeled protein to ~350 μM in NMR buffer containing 10% D2O. KF in a coaxial glass insert was used as chemical shift reference. All data were processed with Bruker Topspin 3.0 and analyzed with NMRViewJ (OneMoon Scientific, Inc). CT-CPMG RD data were analyzed by subtracting effective relaxation rates (*R*_2,eff_) at CPMG field strengths of 25 and 1000 Hz.

### Hydrogen/deuterium exchange mass spectrometry (HDX-MS)

Solution-phase amide HDX experiments were carried out with a fully automated system described previously (Chalmers et al., 2006) with slight modifications. Five μl of apo-Nurr1 LBD was mixed with 20 μL of D_2_O-containing NMR buffer and incubated at 4 °C for a range of time points (0s, 10s, 30s, 60s, 900s or 3,600s). Following exchange, unwanted forward or back exchange was minimized and the protein was denatured with a quench solution (5 M urea, 50 mM TCEP and 1% v/v TFA) at 1:1 ratio to protein. Samples were then passed through an in-house prepared immobilized pepsin column at 50 μL min-1 (0.1% v/v TFA, 15 °C) and the resulting peptides were trapped on a C_18_ trap column (Hypersil Gold, Thermo Fisher). The bound peptides were then gradient-eluted (5-50% CH_3_CN w/v and 0.3% w/v formic acid) across a 1 mm × 50 mm C_18_ HPLC column (Hypersil Gold, Thermo Fisher) for 5 min at 4 °C. The eluted peptides were then analyzed directly using a high resolution Orbitrap mass spectrometer (LTQ Orbitrap XL with ETD, Thermo Fisher). Each HDX experiment was performed in triplicate. To identify peptides, MS/MS experiments were performed with a LTQ Orbitrap mass spectrometer over a 70 min gradient. Product ion spectra were acquired in a data-dependent mode and the five most abundant ions were selected for the product ion analysis. The MS/MS *.raw data files were converted to *.mgf files and then submitted to Mascot (Matrix Science, London, UK) for peptide identification. Peptides with a Mascot score of 20 or greater were included in the peptide set used for HDX detection. The MS/MS Mascot search was also performed against a decoy (reverse) sequence and false positives were ruled out. The MS/MS spectra of all the peptide ions from the Mascot search were further manually inspected and only the unique charged ions with the highest Mascot score were used in estimating the sequence coverage. The intensity weighted average m/z value (centroid) of each peptide isotopic envelope was calculated with the latest version of our in-house developed software, HDX Workbench (Pascal et al., 2012).

### Tryptophan fluorescence spectroscopy

Sixteen final ligand concentrations ranging from 0.23 μM to 300 μM were prepared in ethanol or DMSO (Sigma) from 2 μL of ligand stock (30 mM) or vehicle control (ethanol) spiked to 2 wells per concentration in a 96-well black quartz microplate (Hellma) containing 200 μL of 2.5 μM protein or 25 μM L-tryptophan. After an incubation of 30 min, the plate was read at 23-25 °C in a Tecan Safire II or Molecular Devices Spectramax M5e microplate reader with excitation and emission wavelengths set to 280 nm and 335 nm, respectively; or with an emission scan from 300 to 500 nm at 5 nm increments at the same excitation wavelength. Fluorescence units were converted to percentage fluorescence quenching with respect to vehicle controls (i.e., Nurr1 or L-tryptophan with 1% ethanol or DMSO) and normalized to L-tryptophan data and fit to a one-site specific binding equation with or without Hill slope (determined by F-test in GraphPad Prism) to calculate K_d_ values.

### Molecular dynamics simulations

Generation of MD input files and the production MD runs were performed using AMBER 14 (http://amber-md.org). The ff14SB force field parameters were used for the protein molecule, and ionsjc_tip3p parameters were used for ions (Joung and Cheatham, 2008). The apo-Nurr1 LBD crystal structure (PDB 1OVL) was used to generate coordinate and parameter files within tleap (AmberTools14). The overall charge was neutralized using Na+ ions; TIP3P (Jorgensen et al., 1983) water molecules were added to a minimum thickness of 10 Å surrounding the protein in a truncated octahedron box; and K^+^ and Cl^−^ ions were added to a concentration of 50 mM to be the same as the NMR experiments. The builds were made ready for production runs using a script provided by the Cheatham lab (University of Utah), which involves 4 minimization and 5 equilibration steps in explicit solvent. The equilibration steps are all run at 310K using the Berendsen thermostat to control the temperature. First, minimization is carried out for 1000 steps of steepest descent minimization with strong restraints on the heavy atoms without use of the SHAKE algorithm (Miyamoto and Kollman, 1992). In the second step, molecular dynamics using the NTV ensemble was run for 15 ps with strong restraints on heavy atoms using SHAKE with 1 fs time steps. The third step consists of steepest descent minimization with lower restraints on heavy atoms (no SHAKE), and the fourth and fifth steps repeat the third step with minimal and no restraints on the heavy atoms respectively. The 4 remaining equilibration steps used the NTP ensemble with SHAKE. The first two steps consisted of steadily decreasing restraints on heavy atoms, followed by a third step with steadily decreasing restraints on backbone atoms, for lengths of 5-10 ps and 1 fs time steps. The fourth (final) step removes the restraints entirely and runs 200 ps of MD using the NTP ensemble with 2 fs time steps. The restart file from this final step was then used to start the production MD run. Production MD was performed in the NVT ensemble using the GPU-enabled version pmemd (AMBER14) (Salomon-Ferrer et al., 2013) with hydrogen mass repartitioning (Hopkins et al., 2015) and a 4 fs time step at 310K. Accelerated MD simulations were also performed using AMBER14 using the same minimized restart file with the following flags in the input file as defined in the AMBER manual: iamd = 3, ethreshd=4062, alphad=188.8, ethreshp=-98754, alphap=5511. Trajectory analysis was performed using CPPTRAJ (Roe and Cheatham, 2013) for distance, angle, and DSSP secondary structure measurements (Kabsch and Sander, 1983); and CHIMERA (Pettersen et al., 2004) and VMD (Humphrey et al., 1996) for visualization.

---

**Supplementary Figure S1.**
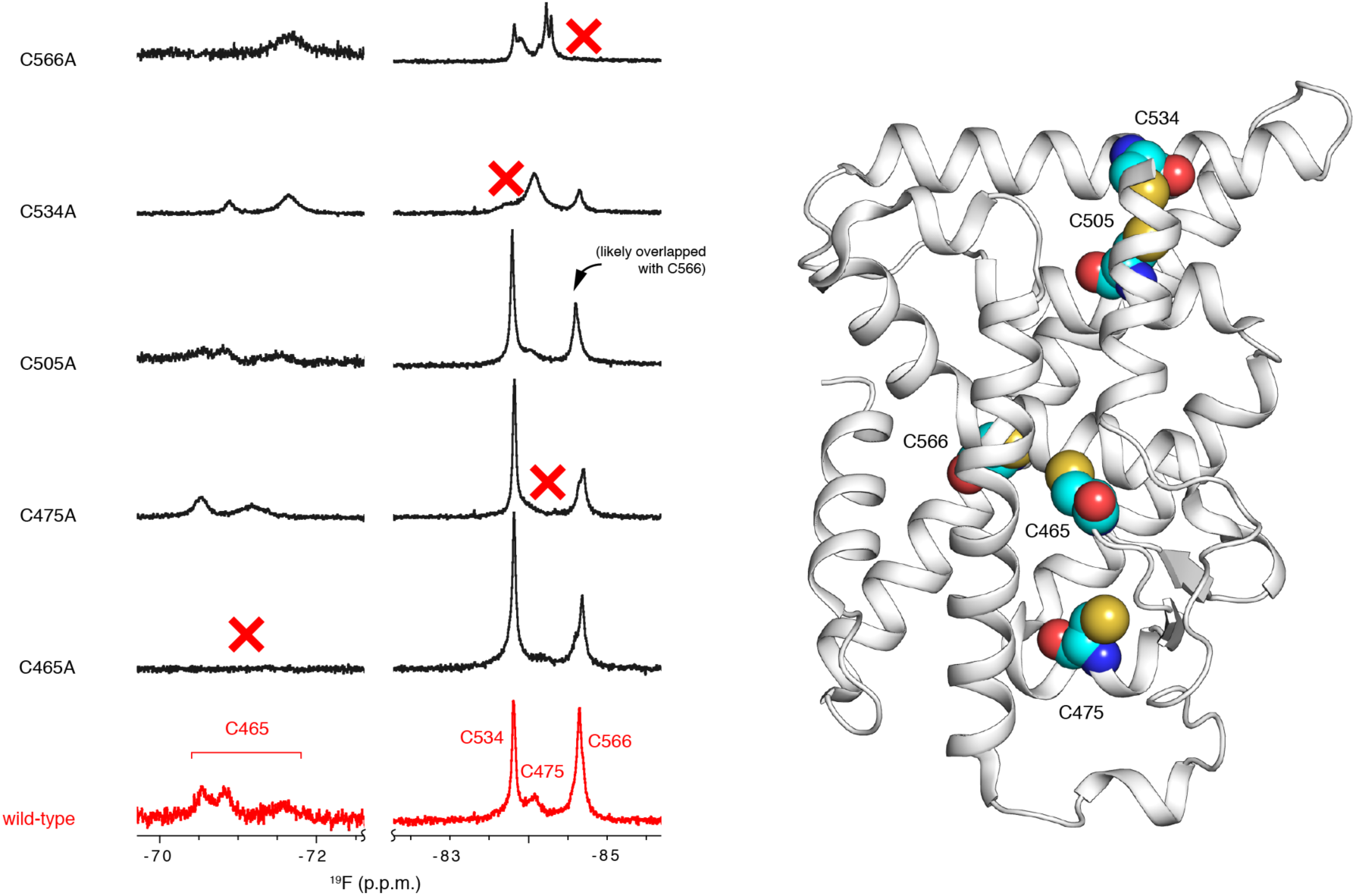
Site-directed mutagenesis was used to individually mutate each of the 5 cysteine residues to alanine in the Nurr1 LBD. Each mutant protein was labeled with BTFA and subjected to ^19^F NMR. Assignment of NMR peaks to cysteine residues was performed via observation of the peak that disappears for each mutant.

**Supplementary Figure S2.**
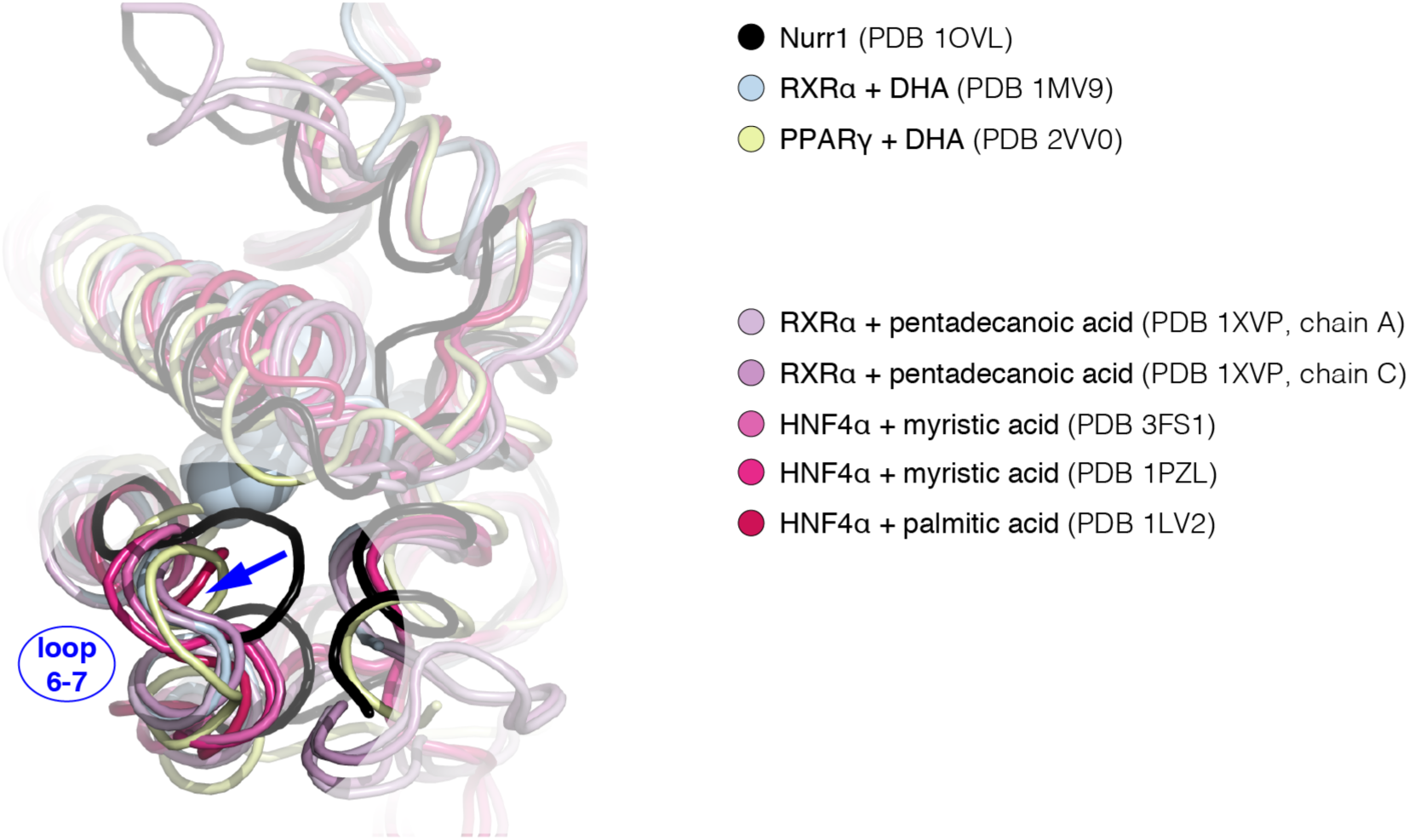
Structural overlay showing that the helix 6–7 loop is in a closed conformation in the apo-Nurr1 LBD crystal structure but an open conformation in crystal structures of other NR LBDs bound to endogenous ligands.

**Supplementary Figure S3.**
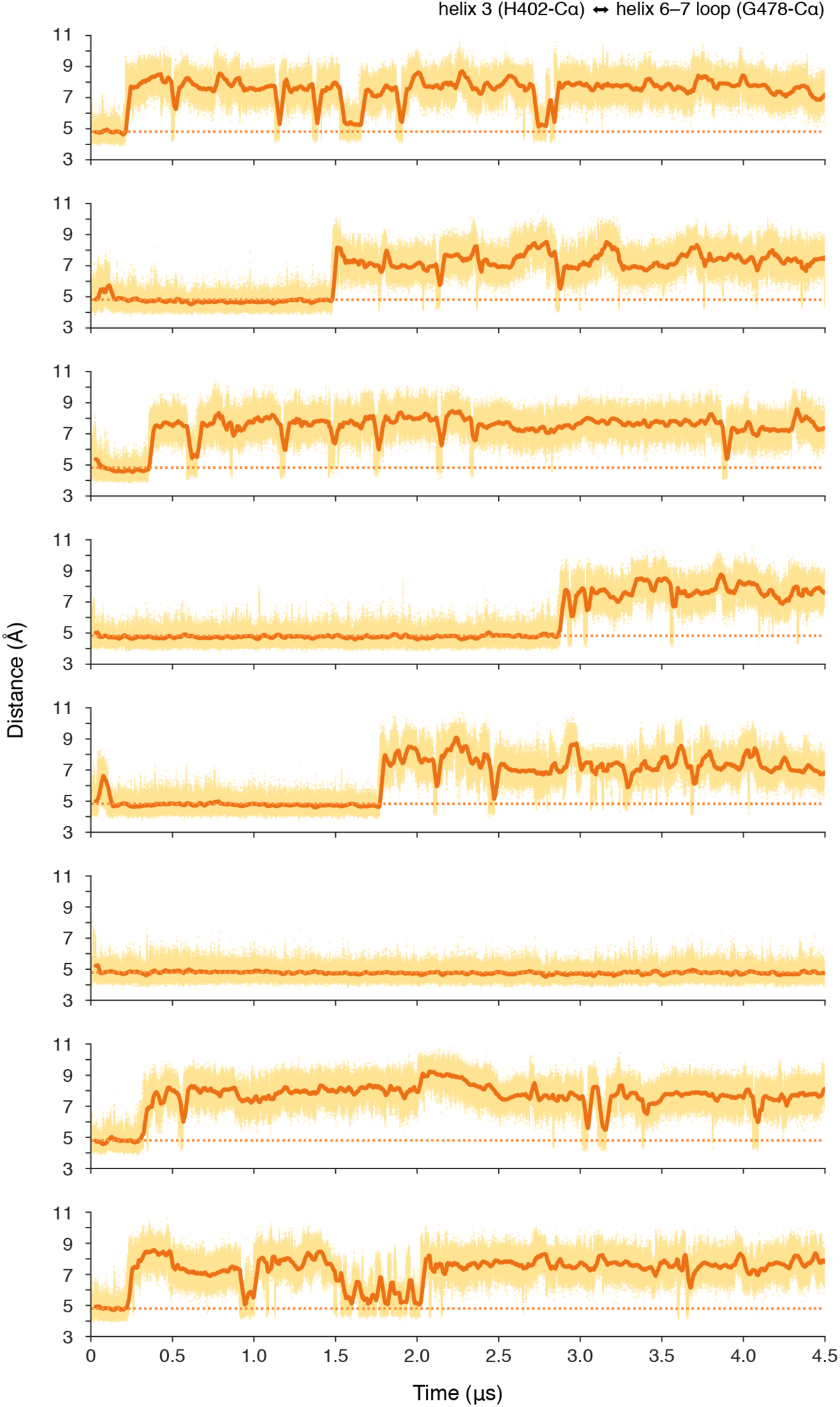
Eight replicate molecular dynamics simulations showing that the helix 6–7 loop (residue G478) moves away from helix 3 (residue H402), transitioning from a closed to open conformation.

